# Modeling the Aging Human Lung: Generation of a Senescent Human Lung Organoid Culture System

**DOI:** 10.1101/2025.02.24.639173

**Authors:** Mohit Aspal, Natalie Pushlar, Melina Melameka, Rachael N. McVicar, Emily Smith, Temiloluwa Ogunyamoju, Matangi Kumar, Jamey D. Marth, Jerold Chun, Evan Y. Snyder, Sandra L. Leibel

**Author notes:** Correspondence &.

## Abstract

**Introduction:** The aging lung enters into a state of irreversible cellular growth arrest characterized by senescence. While senescence is beneficial in preventing oncogenic cell proliferation, it becomes detrimental when persistent, promoting chronic inflammation and fibrosis through the senescence-associated secretory phenotype (SASP). Such senescence-related pathophysiological processes play key roles in lung diseases like chronic obstructive pulmonary disease (COPD) and idiopathic pulmonary fibrosis (IPF). However, few models accurately represent senescence in the human lung.

**Methods:** To generate a human lung senescence *in vitro* model, we first generated a human induced pluripotent stem cell (hiPSC)-derived lung organoid (LO) system which was dissociated into monolayers and air-liquid interface (ALI) cultures to enhance visualization and allow uniform exposure to agents. Cellular senescence was induced using doxorubicin, a DNA-damaging agent. Senescence markers, such as β-galactosidase (β-gal) activity, SASP cytokine production and secretion, cell morphology, proliferative capacity, and barrier integrity were evaluated to validate the senescent phenotype.

**Results:** The doxorubicin-induced senescent hiPSC-derived lung cells demonstrated the hallmark characteristics of cellular senescence, including increased β-gal activity and increased production of the pro-inflammatory SASP cytokine IL-6 and increased secretion of TNF-α. Senescent cells displayed enlarged morphology, decreased proliferation, and reduced wound repair capacity. Barrier integrity was impaired with decreased electrical resistance, and increased permeability, as well as expression of abnormal tight junction proteins and increased fibrosis, all consistent with the senescent lung.

**Conclusion:** Our hiPSC-derived lung cell senescent model reproduces key aspects of human lung senescence and offer an *in vitro* tool for studying age-related lung disease mechanisms and therapeutic interventions. This model has potential applications in exploring the impact of environmental factors (e.g., toxins, infectious pathogens, etc.) on the senescent lung and assessing treatments that could mitigate pathologies associated with pulmonary aging including barrier impairment, inflammation and fibrosis.

## Introduction

Aging is marked by a gradual loss of physiological integrity, resulting in functional decline and increased mortality. According to Centers for Disease Control and Prevention (CDC) data, chronic lower respiratory diseases rank as the 6th leading cause of death in the United States. The lungs, uniquely exposed to external environmental factors, face constant biological and chemical exposures. As lungs age, they are at heightened risk for conditions such as fibrosis, chronic obstructive pulmonary disease (COPD), lung cancer, and infections(1). Key features of aging lungs include enlarged airspaces, increased cellular senescence, decreased elasticity, and increased inflammation(1, 2). Strategies to promote health during aging often focus on clearance of senescent cells with senolytics.

Cellular senescence is a process involving distinct phenotypic changes, such as cell cycle arrest and senescence-associated signaling. Senescence-associated signaling involves key pathways that lead to cellular senescence, notably the p53/p21 and p16^INK4a^/Rb pathways. Activation of these pathways results in cell cycle arrest and the development of the senescence-associated secretory phenotype (SASP) (3, 4). This prevents damaged cells from continuing to propagate and triggers their elimination by the immune system. This process is beneficial when temporary(5); however, prolonged cellular senescence can lead to chronic low-grade inflammation that promotes tissue fibrosis(6). Senescence is classified as either replicative senescence, stress-induced premature senescence (SIPS), or physiological senescence.

Senescent cells release a complex mixture of factors known as the senescence-associated secretory phenotypes (SASP). These include cytokines, chemokines, matrix remodeling proteases, and growth factors(7, 8). These can work in a paracrine fashion and promote the loss of cellular proliferation and tissue deterioration(9) (Fig. 1). Unlike healthy cells, senescent cells do not proliferate and are resistant to both apoptosis and autophagy. In the lungs, senescence of parenchymal and vascular endothelial cells has been implicated in the onset and progression of COPD(10, 11). Lung tissue from patients with chronic pulmonary obstructive disease (COPD) contain a higher proportion of p16-expressing cells and elevated levels of SASP cytokines such as IL-1, IL-6, IL-8, IL-13, and TNF-a(12, 13). Similarly, senescent cells have been identified in the lungs of idiopathic pulmonary fibrosis (IPF) patients, with markers like senescence-associated β-galactosidase (β-gal) and p21 in alveolar epithelial cells(14).

**Figure 1.**
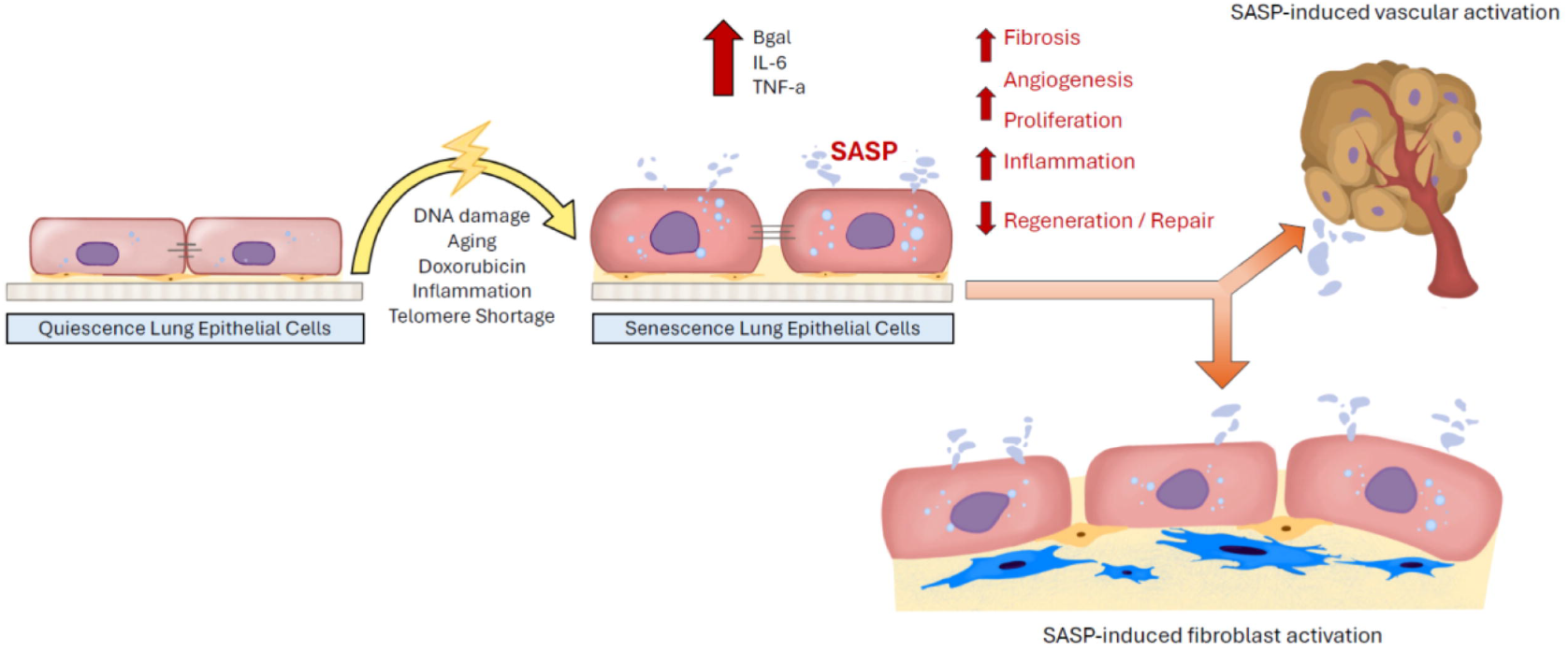
Schematic of senescence induction in lung epithelial cells. Aging-related factors, such as DNA damage, changes quiescent cells into senescent cells which are characterized as being metabolically active but in cell cycle arrest. They secrete SASP, express β-gal, and induce lung fibrosis and vascular activation, resulting in aging-related lung diseases such as COPD and IPF.

Targeted removal of senescent cells through early intervention, such as suicide-gene-mediated ablation, has shown improvements in pulmonary function in bleomycin-injury IPF mouse models(15). Likewise, senolytic drugs have been utilized to selectively clear senescent cells, effectively reducing fibrosis(16).

Animal models have been helpful in replicating some senescence-associated phenotypes, but typically only in specific lung regions, and they do not fully represent the *in vivo* aging human lung(17-20). Human *in vitro* cultures have been developed to study senescence through chemically induced SIPS, utilizing agents like aphidicolin to induce DNA replication stress(21), doxorubicin, to cause DNA damage(22), or pyridostatin as a telomerase inhibitor(23). Senescence induction is validated by observable characteristics such as flattened cell morphology with enlarged nuclei, absence of the proliferation marker ki67, increased lysosomal hydrolases like β-galactosidase(24), and increased SASP production such as IL-6(25). Most human lung *in vitro* models use immortalized pulmonary cell lines or bronchial epithelial cells grown at an air-liquid interface (ALI)(7). While these models can resemble aspects of the senescent human lung, they are limited by the inclusion of only pulmonary epithelial cells. and not the rich diversity of interacting cell types that constitute a fully functioning lung. The lung has many more relevant cell types that develop and function based on their 3D cytoarchitecture and intercellular communication and mutual induction during organogenesis. It is known that SASP produced by the lung epithelium leads to the pathologic activation of mesenchyme resulting in the tissue remodeling, inflammation and fibrosis seen in age-related lung diseases such as COPD and IPF(26).

We have demonstrated the faithfulness and impact of our 3D lung organoid (LO) system(27-30), derived from patient-specific hiPSCs, for accurately modeling pathology throughout the lung and its multiple cell types. In this study, we developed a human *in vitro* model of senescent lung cells using dissociated hiPSC-derived LOs, which contain both epithelial and mesenchymal cells. These organoids were dissociated into monolayers or ALI-cultured epithelium, then exposed to doxorubicin to induce senescence. By creating a model that more accurately represents human lung cellular senescence, we can now gain deeper insights into the biological processes underlying age-related pulmonary diseases and their potential treatments.

## Methods

### Generation of hiPSC-derived lung organoids (LOs), monolayers, and air-liquid interface (ALI) cultures

For hiPSC-LO generation(27-29), we used the ALDA31616 hiPSC cell line which were cultured in feeder free conditions on Matrigel (Corning #354230) coated plates in mTeSR Plus medium (StemCellTech #100-0276). Media was changed daily, and stem cells were passaged once a week using enzyme free dissociation reagent ReLeSR™ (Stem Cell Tech #05872). Cultures were maintained in an undifferentiated state, in a 5% CO_2_ incubator at 37°C. For lung progenitor spheroid generation, hiPSCs were dissociated into single cells with Accutase (StemCellTech # 07920) for 20 minutes, and 2.0 × 10^5^ cells were seeded on Matrigel-coated plates in mTeSR supplemented with 10μM Y-27632 and incubated for 24 hours. Each hiPSC cell line required seeding density optimization resulting in wells that were 50-70% confluent the day after seeding. On day 1, Definitive Endoderm (DE) was induced with DE base medium (RPMI1640+Glutamax, 2% B27 without retinoic acid supplement, 1% HEPES, 50 U/mL penicillin/streptomycin) supplemented with 100 ng/mL human activin A (R&D), 5μM CHIR99021 (Stemgent)(31). On days 2 and 3, cells were cultured in DE media with only 100 ng/mL human activin A supplemented. Anterior Foregut Endoderm (AFE) was induced with serum free base medium (SFBM) supplemented with 10μM SB431542 (R&D) and 2 μM Dorsomorphin (StemGent) on days 4-6(32). SFBM constitutes: 3 parts IMDM:1 part F12, supplemented with 1% B27^-RA^ and 0.5% N2, 50U/mL penicillin/streptomycin, 0.05% BSA, 50 μg/ml L-ascorbic acid, 0.45mM monothioglycerol. On day 7, 2 dimensional (2D) AFE cells were dissociated with Accutase for 10 minutes and 3.0 × 10^5^ cells were passaged as aggregates into 150 μL Matrigel droplets to make Lung Progenitor Cell (LPC) spheroids(29). The Matrigel containing cells were then polymerized for 30-60 minutes in a 37°C incubator and then LPC media (SFBM with 10 ng/mL of BMP4, 0.1 μM of retinoic acid (RA), 3 μM of CHIR99021), supplemented with 10μM Y-27632 the day of passage, was added to submerge the Matrigel droplets. LPC media changed every other day for 9-11 days. Analysis of the surface antigen CPM, or the intracellular marker NKX2-1 was performed at the end of this differentiation period to determine the efficacy of lung progenitor cell differentiation(33, 34). If the LPC spheroids expressed > 50% CPM/NKX2-1, LOs were generated. To generate LOs, LPC spheroids were dissociated using dispase, cold PBS, then TrypLE. LPC spheroids embedded in Matrigel were incubated in 2U/ml dispase for 30 minutes at 37°C. Cold PBS was added to the mixture, transferred to a 15ml conical, then centrifuged at 400 x g for 5 mins. Supernatant was carefully aspirated, leaving 1ml residual solution, 3ml of cold PBS was added to mixture and then centrifuged again at 400 x g for 5 mins. Supernatant was carefully removed and resuspended in 2-3mls of TrypLE Express (Gibco # 12605010) for 12 minutes at 37 °C to keep LPC spheroids as aggregates. Reaction was quenched with 2% fetal bovine serum (FBS) in DMEM/F12 then centrifuged at 400 x g for 5 min. The supernatant was aspirated, and the cell pellet resuspended in 1ml of quenching media supplemented with 10 μM Rock inhibitor (Y-27632) and a cell count was performed. Cells were embedded into Matrigel as aggregates at 5.0 × 10^4^ cells per 200 μL of Matrigel.

To generate 3D human airway LOs, we modified a previously published protocol(35). The medium was composed of serum free basal medium (SFBM) supplemented with 250ng/mL FGF2, 100ng/mL rhFGF10, 50nM dexamethasone (Dex), 100μM 8-Bromoadenosine 3’,5’-cyclic monophosphate sodium salt (Br-cAMP), 100μM 3-Isobutyl-1-methylxanthine (IBMX) and 10 μM ROCK inhibitor (Y-27632). Airway organoid media was changed every other day for 2 weeks.

To generate 2D airway monolayers, human PSC derived LOs were dissociated into single cells and seeded at 20,000 cells per well of a Matrigel coated 96-well plate. LOs embedded in Matrigel were incubated in 2U/ml dispase for 30 minutes at 37°C. Cold PBS was added to the mixture and then centrifuged at 400 x g for 5 mins. Supernatant was carefully aspirated, leaving 1ml residual solution, 3ml of cold PBS was added to mixture and then centrifuged again at 400 x g for 5 mins. Supernatant was carefully removed and resuspended in 2-3mls of TrypLE Express (Gibco # 12605010) for 20 minutes at 37 °C to obtain single cell solution. Reaction was quenched with 2% FBS in DMEM/F12 then centrifuged at 400 x g for 5 min. The supernatant was aspirated, and the cell pellet resuspended in 1ml of quenching media supplemented with 10 μM Rock inhibitor (Y-27632). Cell count was performed, and the respective volume of cells were resuspended in Pneumacult Ex+ media (StemCellTech) into 96 well plates at 100 μL per well as monolayers.

To generate ALI-cultures, hiPSC derived LOs were dissociated into single cells (as described above) and seeded at 3.0 × 10^5 cells onto the apical side of a Matrigel coated 3.0 μm pore polyester transwell insert (see above for dissociation protocol). Once the cells were confluent, the apical media was removed, and the basolateral chamber was changed with PneumaCult ALI media (StemCellTech) every 48 hours.

### Senescence induction via Doxorubicin

HiPSC-derived LOs were dissociated as described above, plated as 2D monolayers, and treated with doxorubicin for 24 hours at 37°C at various concentrations (0.0, 0.3, 0.5, 1.0, 2.0, or 5.0 μM doxorubicin diluted in LO media). After 24 hours, doxorubicin was aspirated, and cells were returned to Doxorubicin-free LO media if submerged or were air-lifted if in ALI-culture, for 7 days. After 7 days, the cells were fixed for immunofluorescence staining or stained for β-gal expression. Supernatants from senescent cultures were collected and examined for the secretion of SASP. Control cultures treated with DMSO were harvested along a similar timeline.

### Immunofluorescence

2D monolayers and ALI-cultures of hiPSC-derived LOs were fixed in 4% paraformaldehyde (PFA) for 15-60 minutes and then washed twice with PBS. For immunofluorescence, cells were blocked with 5% v/v donkey serum and 5% w/v bovine serum albumin (BSA) with 0.1% Triton X-100 for 1 hour at room temperature (RT). Cells were then incubated in primary antibody diluted in blocking buffer overnight at 4°C. The next day, cells were stained with secondary antibodies (ThermoFisher) at RT for 1 hour. Control slides were stained with secondary antibodies only. Images were taken on a Keyence BZ-X fluorescent microscope and processed and analyzed using ImageJ software.

### Antibodies

We used IL-6 (Invitrogen P620), β-gal (Invitrogen C10840), Ki67 (Santa Cruz Biotech sc-23900) Vimentin (Santa Cruz Biotech sc-6260), and SMA (Invitrogen, 50-9760-80).

### Flow Cytometry

Flow cytometry staining was performed according to manufacturer’s recommendations for the Invitrogen CellEvent Senescence Green Flow Cytometry Assay Kit (Invitrogen C10840). In brief, doxorubicin-induced senescent hiPSC lung monolayers were single-cell dissociated, washed with PBS, then incubated in 5% Hoechst in Hank’s balanced salt solution (HBSS) for 15 min at RT. Cells were centrifuged at 200 x g for 5 mins, washed with PBS, and resuspended in Fixation Solution (2% Paraformaldehyde in PBS) for 10 min at RT. Cells were centrifuged at 400 x g for 5 mins, washed with 1% BSA in PBS, and incubated with primary β-gal antibody (1:1000 CellEvent Senescence Green Probe in CellEvent Senescence Buffer) for 1 hour at 37°C protected from light. After washing again twice with 1% BSA in PBS, cells were resuspended in 300 μL 1% BSA before flow cytometry analysis. Flow cytometry was performed using unstained hiPSC-LOs as a negative control and gates were set to exclude the dead cells (DAPI stained). Flow cytometry was done on the BD LSRFortessa Cell Analyzer.

### MTT [3-(4,5-dimethylthiazol-2-yl)-2,5-diphenyltetrazolium bromide] Colorimetric Assay

HiPSC LOs were plated as 2D monolayers on a Matrigel coated Costar 96-well transparent polystyrene plate (3.0 × 10^4 cells/well), induced with doxorubicin for 24 hours, and placed in DMEM + F12 media without phenol red, the night before the MTT assay. 10 μL of MTT substrate was added to each well and incubated for 3 hours at 37°C protected from light. After 3 hours, 100 μL of SDS is added to each well and left on a plate shaker overnight at RT protected from light before reading absorbances at 570nm using the Infinite 200Pro plate reader. Nine absorbance reads were measured for technical replicates for each well.

### Scratch Assay

HiPSC LOs were plated as 2D monolayers on an Matrigel coated IncuCyte Imagelock 96-well plate (Sartorius BA-04857), induced with doxorubicin for 24 hours, and washed once with 1X PBS before the scratch assay. Uniform linear scratches were created in each well using the IncuCyte 96-well Woundmaker Tool (Sartorius Cat# 4563). Wells were washed twice with 1X PBS, 200 μL airway LO maturation media was added, and then the plate was placed in an IncuCyte Live-Cell Analysis System at 37°C to measure the progress of wound closure every hour for 24 hours. The sizes of the scratches were measured using the IncuCyte Scratch Wound Analysis software module. 3 images were acquired per well, and the average wound width (μm) per timepoint was evaluated using the Scratch Wound module.

### TNF-α ELISA

ELISA assay for TNF-α was performed using the Invitrogen Human TNF-α Sandwich ELISA Kit (Invitrogen BMS223-4). After the supernatant of infected cells was collected, 50 μL of sample dilutant was added to 50 μL supernatant samples in ELISA strip wells included with kit and incubated with 50 μL Biotin-Conjugate for 2 hours at RT protected from light. Wells were washed four times with 1X wash buffer and incubated with 100 μL Strep-HRP for 1 hour at RT protected from light. Wells were washed again four times with 1X wash buffer, incubated with 100 μL TMB substrate solution, and stopped with 100 μL Stop Solution after 10 min.

Absorbances were read for each well at 450nm using the Infinite 200Pro plate reader. ELISA was performed using reconstituted TNF-α standards (included with kit) as positive controls. Blank well with only sample dilutant was used to calculate corrected absorbance values for all standard and sample wells.

### Transepithelial electrical resistance (TEER)

TEER was performed using an EVOM2 with STX2/chopstick electrodes. One electrode was positioned in the apical chamber and the other in the basolateral chamber(36). A low-frequency AC current (typically 12.5 Hz) was applied and the voltage was measured across the cell layer. The TEER value was calculated by subtracting the blank resistance (membrane without cells) from the total resistance and multiplying by the effective membrane area (TEER = (R_total - R_blank) × Area). The final TEER value was expressed in Ω·cm^2^. Measurements were repeated 3 times per membrane (technical replicates) and in 3 different transwells (biological replicates).

### Dextran-FITC permeability assay

ALI-cultured cells were analyzed after 7 days post doxorubicin treatment. Prior to the MTT assay(37), the culture medium was replaced with fresh, phenol red-free medium in both the apical and basolateral chambers. Fluorescein isothiocyanate (FITC)-conjugated dextran (4 kDa) was added to the apical chamber at a final concentration of 1 mg/mL. The inserts were then incubated with dextran at 37°C in 5% CO2 for 4 hours. Then 100 μL of media was collected from the basolateral chamber. The fluorescence intensity of the collected samples was measured using a microplate reader at excitation and emission wavelengths of 485 nm and 528 nm, respectively. The fluorescent reading and respective dextran concentration was calculated via a standard curve with known dextran concentrations. Undiluted dextran-FITC is 25 mg/ml, dilutions at 1:200, 1:100, 1:66, 1:50, 1:33, 1:25 and a PBS negative control were evaluated in triplicate. The assay was performed in triplicate for each experimental condition.

### Statistical Analysis

Results were represented as mean ± standard error (SE) of the mean or standard error as indicated. The comparison between two groups was performed by unpaired non-parametric Mann–Whitney test or t-test with Welch’s correction using GraphPad Prism (GraphPad Software, Inc., La Jolla, CA). Correlation analysis between indicated parameters was also conducted by non-parametric Spearman test. Normalized values ranged between 0 and 1. Ordinary one-way ANOVA tests were performed without pairing and assuming Gaussian distribution. Multiple comparison tests were performed under each ANOVA to compare significance of data between different experimental conditions. A p-value of ≤ 0.05 was considered statistically significant.

## Results

### Development of hiPSC-derived senescent models from LO cultures

HiPSC-derived LOs were dissociated, plated as monolayers, and treated with escalating concentrations of doxorubicin for 24 hours using a modified protocol(38). Lysosomal hydrolase β-galactosidase (β-gal) was subsequently analyzed via flow cytometry (Fig. 2A). Control cells exhibited 1.8% β-gal expression, which increased proportionally with higher doxorubicin concentrations, reaching a peak at 1.0 μM (Fig. 2B). Doses as high as 5.0 μM led to increased cell death and a reduction in β-gal expression (Fig. 2B). We further examined β-gal expression after doxorubicin removal over time and found that β-gal-positive cells rose to 73% by day 7, then decreased to 63% by day 10 (data not shown).

**Figure 2.**
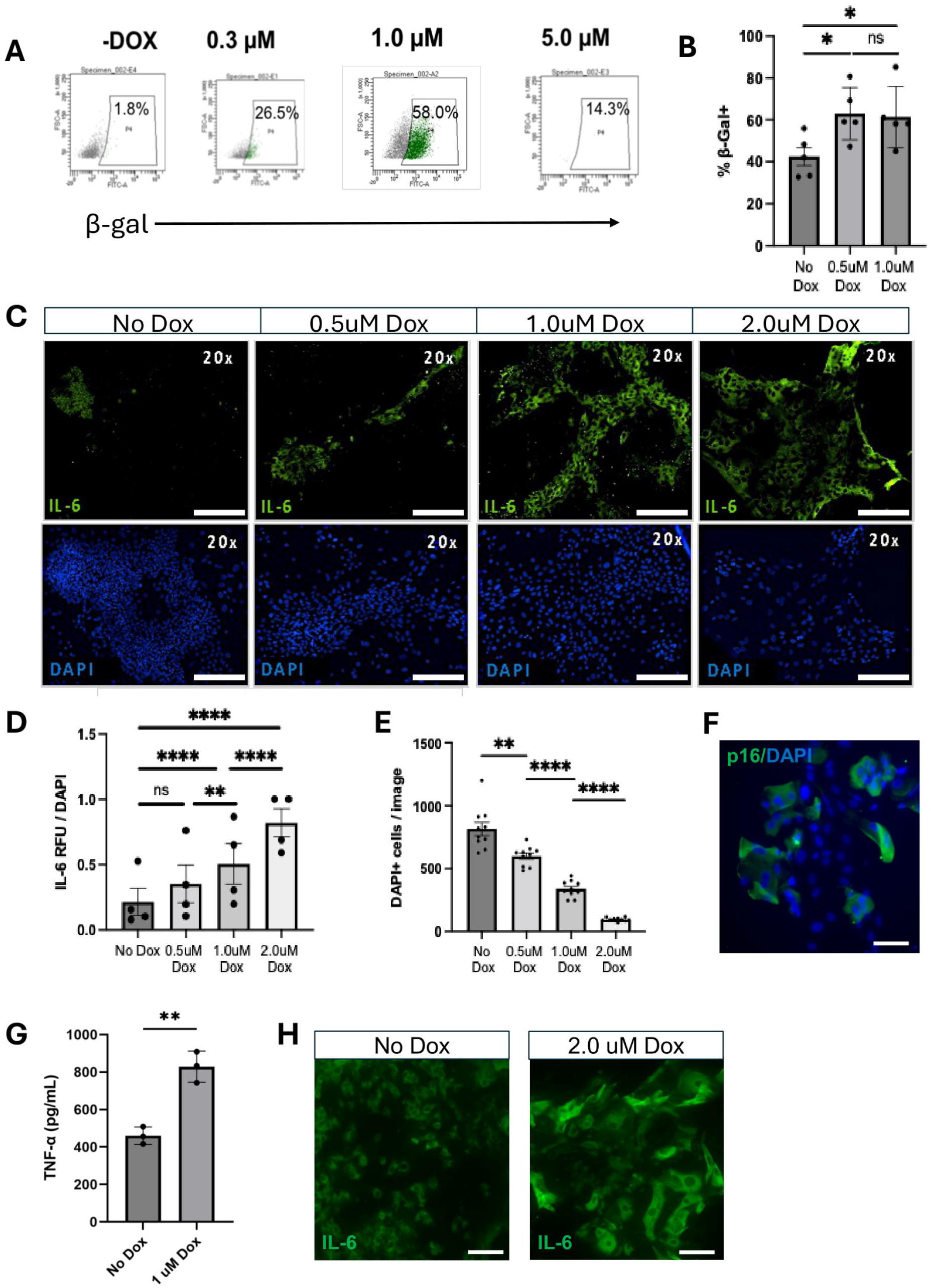
Validation of Doxorubicin-induced senescence in hiPSC lung organoid (LO)-derived monolayers. **A**) Flow cytometry of dissociated LO cells for β-gal 7 days post treatment with doxorubicin at various concentrations. Peak is achieved at 1.0 μM with a diminution if escalates as high as 5.0 μM. **B**) Bar graph quantifying the number of cells expressing β-gal from the flow cytometry assay. N= 5 biological replicates. Statistics by t-test. Error bars +/-standard error of the mean (SEM). p-value < 0.05 is significant; p-value > 0.05 is not significant (ns). **C**) Representative immunofluorescent images for IL-6 (green signal) one week post doxorubicin treatment at various concentrations. Images 20x, scale bar 200 μM. **D**) Bar graph quantifying the mean relative fluorescent intensity of IL-6. Relative fluorescent units (RFU) normalized per number of DAPI+ cells per image. N = 4 biological replicates. 7 image replicates taken per differentiation. Statistics by one-way ANOVA. Error bars +/-SEM. P-value <0.05 * <0.01 ** <0.001 *** <0.0001 ****. **E**) Bar graph quantifying the number of DAPI + cells as a marker of cell number 7 days post-Doxorubicin treatment at escalating concentrations. Images taken at 20x. N = 10 biological replicates. Statistics by t-test. Error bars +/-SEM. P-value <0.01 ** <0.0001 ****. **F**) Representative immunofluorescent image for p16 (green signal) one week post 1.0 μM doxorubicin treatment. Images 20x, scale bar 200 μM. G) Bar graph quantifying the release of TNF-α into the cell media. Statistics by t-test. Error bars +/-SEM. P-value <0.01 ** **H**) Morphology of the monolayer cells after doxorubicin exposure and staining for IL-6. Cells exposed to 2 μM of doxorubicin were larger and flatter than the control cells.

SASP production was assessed by staining lung cell monolayers for the representative surrogate molecule intracellular IL-6 (Fig. 2C, D). IL-6 expression showed a dose-dependent increase, with relative fluorescence intensities rising in proportion to the number of DAPI + cells (Fig. 2D). The number of DAPI+ cells declined with higher doxorubicin concentrations despite identical seeding densities across monolayers prior to treatment (Fig. 2E). The optimal doxorubicin dose based on IL-6 secretion and cell number would appear to be 1.0 μM. We further measured SASP secretion in response to doxorubicin by collecting culture supernatants after 7 days and quantified TNF-α levels. Cells exposed to 1.0 μM doxorubicin secreted nearly twice the TNF-α of controls (Fig. 2F). Therefore, a 1.0 μM doxorubicin dose range appeared to be appropriate by this metric, as well.

Morphologically, IL-6+ cells exposed to doxorubicin displayed an enlarged, flattened appearance compared to controls, a morphology consistent with senescent cells (Fig. 2G).

Therefore, treatment with 1.0 μM doxorubicin for 24 hours, followed by 7 days in standard media, was sufficient to induce markers of senescence in hiPSC-derived lung monolayers.

### Senescent lung cells show a reduction in time to wound healing and proliferation

To assess whether the model reflects the reduced wound-healing capacity characteristic of aging, we conducted a scratch assay on hiPSC-derived lung monolayers (Fig. 3A). When monolayers reached 70–80% confluence, doxorubicin was applied at various concentrations for 24 hours. Cells were then maintained in drug-free media for 7 days before performing the scratch assay. Wound recovery (distance from scratch edge) was evaluated by brightfield imaging at hourly intervals. After 24 hours, untreated cells showed significantly smaller wound widths compared to those exposed to 1.0 μM doxorubicin (Figs. 3B–D).

**Figure 3.**
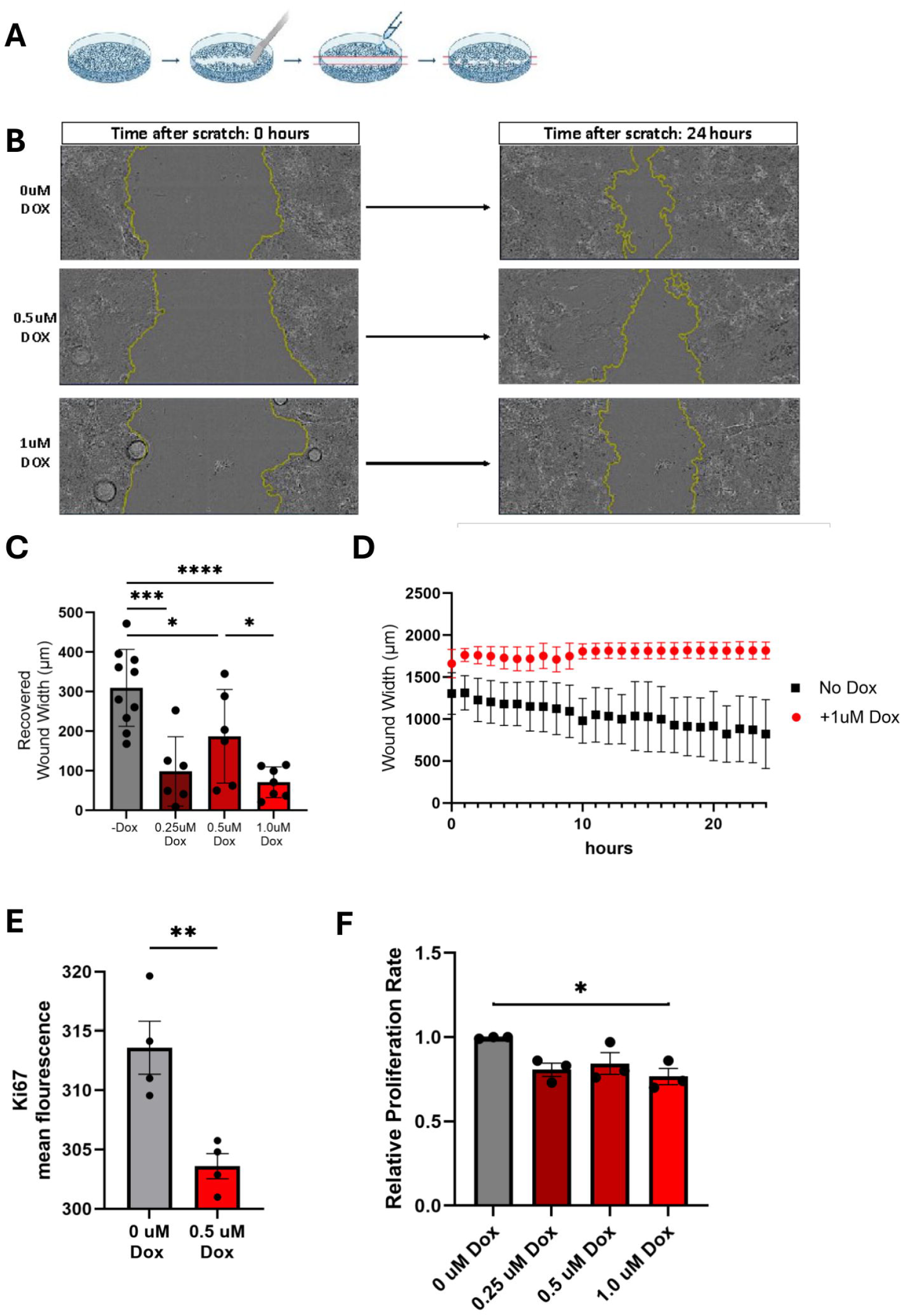
Doxorubicin-induced senescence decreases lung epithelium repair function *in vitro* as measured by poorer performance on the scratch wound closure assay and diminished markers of cell proliferation (Ki67) **A)** Schematic of the scratch assay. **B**) Representative brightfield images of the scratch assay. Yellow lines outline the edge of the wound. **Panels on the left** show the wound closure right after the scratch and the **right panels** show wound closure after 24 hours. The **top panels** represent the control cells, the **middle panels** represent the 0.5 μM doxorubicin exposed cells, and the **bottom panels** represent the 1.0 μM doxorubicin exposed cells. **C**) Bar graph quantifying the total distance of wound recovery post-scratch within a 24 hour period by doxorubicin dose. N= 3-5 biological well replicates, 2 images per well. Error bars +/-standard deviation. Statistics by t-test. P-value <0.05^*^ <0.01^**^ <0.001^***^ <0.001^****^. **D**) Dot plot quantifying the average wound width over 24 hours post-scratch in control and doxorubicin exposed cells. N= 6 biological replicates, 3 images per well. Error bars +/-standard deviation (SD) **E**). Bar graph of the respective mean fluorescent intensity of Ki67 per image. N=4 biological replicates. Error bars +/-SEM. Statistic by t-test. P-value <0.01^**^. **F**) Bar graph illustrating the relative proliferation rate of hiPSC-derived lung monolayers exposed to escalating doxorubicin concentrations via a MTT proliferation assay. Proliferation was assessed over a 24-hour period. Tetrazolium MTT absorbance normalized. 3 experimental replicates, 2 well replicates, 3 reading replicates per well. Error bars +/-SEM. Statistics by one-way ANOVA. P-value <0.05 *

We further assessed proliferation by staining for the marker Ki67, which revealed markedly higher expression in control cells compared to doxorubicin-treated cells (Fig. 3E). To evaluate cell viability, an MTT assay was conducted on both control and doxorubicin-treated cells at different concentrations. Results indicated that doxorubicin induction resulted in significantly decreased mitochondrial dehydrogenase activity by MTT, signifying decreased proliferation compared to controls (Fig. 3F).

Taken together, these findings demonstrate that doxorubicin-induced senescence in hiPSC-derived LO monolayers impairs wound healing and cell proliferation, mimicking aspects of aging-related functional decline.

### Barrier integrity is impaired in senescence

Barrier integrity is essential for maintaining lung health, serving as the first line of defense against environmental insults. With barrier impairment, the risk of injury rises. To assess whether senescence induction compromised barrier integrity, we measured transepithelial electrical resistance (TEER) on day 7 post doxorubicin-induction of hiPSC-derived ALI cultures (Fig. 4A). Doxorubicin-treated cells exhibited significantly lower resistance compared to controls (Fig. 4B). This finding was corroborated by a Dextran-FITC assay, which revealed markedly elevated permeability in doxorubicin-treated cells (Fig. 4C).

**Figure 4.**
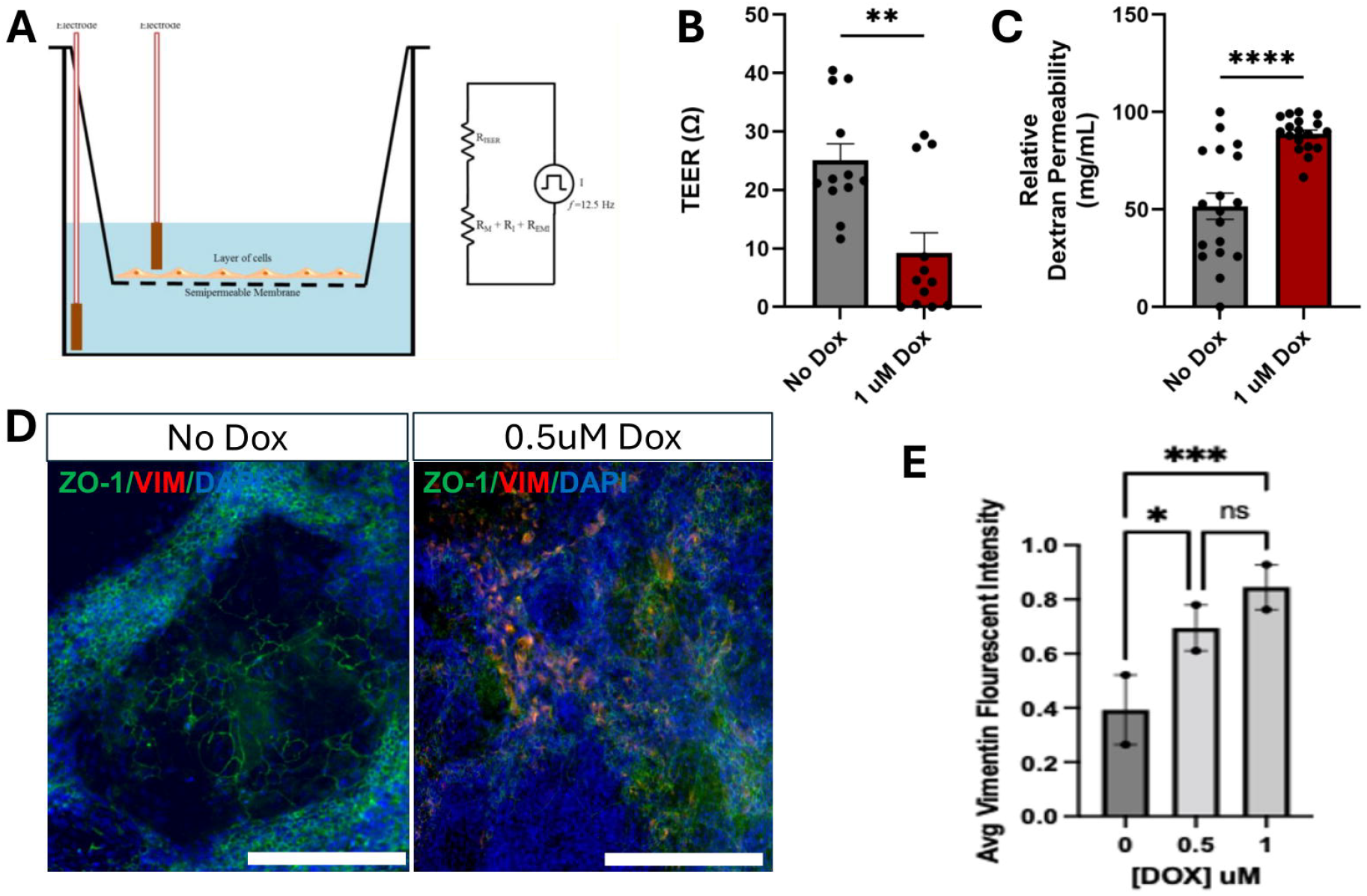
Doxorubicin-induced senescence impairs the lung epithelial barrier. **A**) Schematic of the TEER system in a transwell with apical cells. **B**) Bar graph quantifying the resistance of ALI-cultured hiPSC-derived LOs with and without doxorubicin. Less resistance indicates poorer barrier function. Three experimental replicates, 3 transwell replicates. Error bars +/-SEM. Statistics by t-test. P-value <0.05^*^ <0.01^**^. **C**) Bar graph quantifying the permeability of ALI-cultured hiPSC-derived LOs with and without doxorubicin. The fluorescence intensity of the collected basolateral and apical samples was measured using a microplate reader at excitation and emission wavelengths of 485. A standard curve was also analyzed using varying concentrations of Dextran-FITC. In this assay, increased dextran indicates worse barrier integrity because of increased permeability. The assay was performed in triplicate for each experimental condition. Error bars +/-SEM. Statistics by t-test. P-value <0.001^****^. **D**) Immunofluorescence of the epithelial tight junction protein ZO1 and the mesenchymal cell maker vimentin in doxorubicin treated hiPSC LO-derived monolayers. DAPI is a nuclear stain. In this assay, a lower expression of ZO1 indicates reduced barrier function and an increase in vimentin represents increased epithelial-to-mesenchymal transition due to cellular senescence. Images 10x, scale bar 200 μM. **E**) Bar graph quantifying the fluorescence intensity of vimentin in control cells vs doxorubicin exposed cells. Greater vimentin is a precursor to greater fibrosis. Error bars +/-SEM. Statistics by one-way ANOVA. P-value <0.05^*^ <0.001^***^.

To investigate barrier protein integrity, we immunostained for ZO1 and observed intact expression in control cells, while doxorubicin-treated cultures showed reduced ZO1 expression (Fig. 4D). Given that SASP can promote paracrine signaling and induce fibroblast activation, we further immunostained the doxorubicin-treated cultures with vimentin. The fluorescence intensity significantly increased compared to control cells, suggesting fibroblast activation, the precursor to fibrosis.

Taken together, these results demonstrate that senescence-induced cells exhibit impaired barrier integrity, shown by reduced transepithelial electrical resistance, increased permeability, loss of tight junction protein expression, and elevated fibroblast markers.

## Discussion

Few models of the lung to date are equipped to accurately represent senescence in a complete representation of the multi-cell type human lung, including not only pulmonary epithelium but also mesenchyme. We believe that in a number of prior publications we have demonstrated the faithfulness and impact of our hiPSC-derived LO system for modeling pathology throughout the lung and its multiple cell types.

Our results demonstrate that our hiPSC-derived LO-derived monolayer system, when treated with doxorubicin, exhibit key markers of cellular senescence. Treatment with doxorubicin resulted in dose-dependent increases in senescence-associated β-gal expression, particularly at the 1.0 μM concentration, suggesting that this dose is effective in promoting senescence without causing excessive cytotoxicity. Interestingly, while doses of doxorubicin (as high as 5.0 μM) initially increased β-gal expression, it also led to increased cell death and a subsequent reduction in β-gal+ cells, highlighting the importance of selecting an optimal doxorubicin concentration to avoid excessive cell loss when modeling senescence.

Over time, after removing doxorubicin, we observed that the percentage of β-gal+ cells peaked at 73% on day 7 and declined slightly to 63% by day 10, indicating that senescent cells persist but may undergo turnover or loss over prolonged culture. This aligns with previous studies showing that senescent markers may fluctuate as cells adapt to *in vitro* conditions post-treatment. A significant dose-dependent response of IL-6 expression was observed at increasing doxorubicin concentrations. The induction of the SASP was also confirmed by the secretion of TNF-α, which increased significantly in response to 1.0 μM doxorubicin. The dose-response relationship observed in IL-6 expression, as well as the elevated TNF-α levels in the culture supernatant, confirms that our model can effectively induce SASP production, a hallmark of cellular senescence. These findings are particularly relevant, as the release of pro-inflammatory SASP factors contributes to chronic inflammation and tissue damage, and fibrosis, which are known contributors to age-related lung disease progression.

Taking together the data in Fig. 2 (maximal β-gal and SASP expression while avoiding cytotoxicity), the optimal dose of DOX would appear to be 1.0 μM. This dose appears to be optimal for inducing the other hallmarks of aging described below.

The morphological changes observed in IL-6 + cells, characterized by enlarged and flattened cells, further support the induction of senescence in these cells. These changes are consistent with the known features of senescent cells, which typically exhibit increased cell size and altered shape. Taken together, these findings indicate that a 24-hour treatment with 1.0 μM doxorubicin followed by 7 days of recovery in normal media is sufficient to induce a stable senescent phenotype in hiPSC-derived lung monolayers, evidenced by elevated β-gal activity, SASP production, and morphological changes.

Our study demonstrates that a doxorubicin-treated system can also reflect functional consequences of lung cellular senescence. Wound healing and cellular proliferation were impaired, modeling age-related declines in lung regenerative capacity. The scratch assay results show that doxorubicin-treated cells exhibit slower wound closure compared to untreated controls, which is consistent with the diminished repair capacity observed in aging lung tissue.. This impairment aligns with the accumulation of senescent cells, which exhibit cell cycle arrest and reduced proliferative potential(39). These characteristics hinder the ability of epithelial cells to regenerate and close wounds efficiently. Additionally, the SASP, characterized by the release of pro-inflammatory cytokines, proteases, and growth factors, may contribute to a pro-inflammatory environment that disrupts normal tissue repair, leading to prolonged inflammation and tissue remodeling, ultimately impairing repair(40, 41).

However, we acknowledge that senescence can also have a beneficial role in certain contexts, such as promoting the resolution of fibrosis or enhancing the recruitment of regenerative cells during early repair. The interplay between these opposing effects likely depends on factors such as the duration of senescence, the specific cell types involved, and the balance of SASP components(42).

Future studies could focus on modulating the senescence response, such as selectively targeting the SASP or clearing senescent cells, to better understand how these approaches might restore or even enhance wound repair capacity in this model.

The reduced expression of Ki67, a marker of cell proliferation, in doxorubicin-treated cells further supports the establishment of a senescent phenotype in this model. These findings align with the known characteristics of senescent cells, which undergo cell cycle arrest and display decreased proliferative activity, impairing tissue repair. Additionally, the MTT assay showed reduced cell viability in doxorubicin-treated cells, reinforcing the notion that senescence induction affects cellular function and viability in a dose-dependent manner.

Maintaining barrier integrity is essential for lung health, as it serves as the primary defense against environmental exposures. In our study, we demonstrated that senescence induction, via doxorubicin treatment, significantly impairs barrier function. Reduced TEER in doxorubicin-treated cells indicates weakened barrier integrity compared to untreated controls. This finding was further supported by the Dextran-FITC assay, which showed increased permeability in senescent cells. This increased permeability in senescent cells suggests formation of a compromised barrier that may heighten susceptibility to injury, pathogen incursion, and environmental insult, conditions often associated with aging and age-related lung disease.

The reduction in tight junction protein ZO1 expression in doxorubicin-treated cultures illustrates the structural impact of senescence on epithelial barrier components. Tight junctions are critical for maintaining selective permeability and regulating paracellular transport; their reduction in senescent cells is a key indicator of compromised epithelial function. Loss of ZO1 expression aligns with prior observations in aging lung models, where tight junction disruption is linked to heightened vulnerability to pathogens and environmental toxins. The observed decline in tight junction integrity due to senescence reinforces the connection between cellular aging and impaired lung barrier function.

Additionally, we observed paracrine effects of senescent cells on the surrounding cell types, particularly fibroblasts, as indicated by increased vimentin expression in doxorubicin-treated cultures. Elevated vimentin levels reflect fibroblast activation, suggesting that senescent cells may contribute to a pro-fibrotic environment by releasing SASP that signal fibroblast activation. This fibroblast response can drive fibrosis and tissue remodeling, processes often seen in chronic lung conditions such as COPD and IPF. By inducing fibroblast activation, senescent cells can contribute to the structural and functional decline associated with age-related lung disease.

One limitation of our doxorubicin-induced senescence model is that iPSC-derived lung organoids were dissociated into ALI cultures and monolayers rather than inducing senescence within intact organoids. This may limit its applicability as a full representation of the aged lung. However, many of our functional assays (eg. wound healing, barrier function) cannot be performed in a 3D system. Therefore, we prioritized enhancing the reproducibility of the model by dissociating lung organoids into ALI cultures and monolayers to ensure better visualization, controlled doxorubicin exposure, and more accurate functional assessments. Incorporating 3D lung organoid-based senescence models in future studies would enhance the relevance of this approach for aging and lung disease research.

Another limitation is the absence of immune cells, particularly macrophages and T cells, which play a critical role in senescence surveillance and clearance. The lack of immune interactions may limit our ability to fully capture the *in vivo* dynamics of senescent cell persistence and removal. Future studies incorporating co-culture systems with immune cells may help address this limitation and enhance the model’s translational value.

Additionally, the long-term stability of the senescent phenotype remains to be explored, as indicated by the decline in β-gal activity by day 10. Senescent cells may undergo apoptosis, immune-mediated clearance, or shifts in SASP dynamics, which could influence the functional relevance of the model over time.

## Conclusion

This model presents an effective approach for studying human lung cellular senescence *in vitro* and provides a foundation for future investigations into the mechanisms of age-related lung diseases and for evaluating senolytic therapies or interventions targeting the senescence-associated secretory phenotype. Our results suggest that hiPSC-derived lung monolayers could serve as a valuable *in vitro* platform for examining the impact of various interventions on senescent lung cells particularly in combination with defined disease susceptibility genotypes of hiPSC donors, thereby contributing to the development of novel models and treatments for targeting senescence in chronic lung disease.

## Funding Information

S.L.L. and E.Y.S are supported by grants from the SENS Research Foundation and the California Institute of Regenerative Medicine (DISC2COVID19-12022)

E.Y.S. is supported by the Richmond Family Fund and Sanford Burnham Prebys Medical Discovery Institute

J.D.M. is supported by NIH grants HL131474, HL158677, AI151371, and DK048247.

J.C. is supported by NIH R01 AG065541 and R01 AG071465

## Author Contributions

SLL, MA, NP, RNM designed research; MA, NP, MM, RNM, ES, TO, MK, CM, SLL performed research; MA, NP, RNM, ES, TO, MK, CM, SLL analyzed data; SLL, MA, RNM, EM wrote the paper; MA, NP, RNM, ES, MK, JDM, JC, EYS, SLL edited the paper; SLL, JDM, JC, EYS contributed reagents, tools, and funding.

## Competing Interests

Dr. Chun has an employment relationship with Neurocrine Biosciences, Inc., a company that may potentially benefit from the research results. Dr. Chun’s relationship with Neurocrine Biosciences, Inc. has been reviewed and approved by Sanford Burnham Prebys Medical Discovery Institute in accordance with its Conflict of Interest Policies. The current work has no relationship nor overlap with work at Neurocrine.

## Notes

### Competing Interest Statement

The authors have declared no competing interest.

